# Non-random organization of flux control mechanisms in yeast central metabolic pathways

**DOI:** 10.1101/2021.12.15.472747

**Authors:** Rosemary Yu, Egor Vorontsov, Carina Sihlbom, Jens Nielsen

**Author notes:** **Corresponding author:** Jens Nielsen.

## Abstract

Metabolic flux can be regulated by a variety of different mechanisms, but the organization of these mechanisms within the metabolic network has remained unknown. Here we test the hypothesis that flux control mechanisms are not distributed randomly in the metabolic network, but rather organized according to pathway. Combining proteomics, phosphoproteomics, and metabolic modeling, we report the largest collection of flux-enzyme-phosphoenzyme relationships to date in *Saccharomyces cerevisiae.* In support of the hypothesis, we show that (i) amino acid metabolic pathways are predominantly regulated by enzyme abundance stemming from transcriptional regulation; (ii) upper glycolysis and associated pathways, by inactivating enzyme phosphorylation; (iii) lower glycolysis and associated pathways, by activating enzyme phosphorylation; and (iv) glycolipid/glycophospholipid pathways, by a combination of enzyme phosphorylation and metabolic compartmentalization. We delineate the evolutionary history for the observed organization of flux control mechanisms in yeast central metabolic pathways, furthering our understanding of the regulation of metabolism and its evolution.

## Introduction

Metabolism is a core cellular function and is subject to extensive regulation. Several molecular mechanisms have been identified to regulate metabolic flux, but whether these flux control mechanisms are randomly distributed in the metabolic network, or organized according to a certain structure, has not been examined. Moreover, if the distribution is non-random, how the organizational structure has evolved is unknown.

The metabolic network can be divided into a number of inter-connected pathways, each describing a chain or a cycle of reactions wherein the product of one reaction act as the substrate for the next. Pathways in central metabolism, in particular, consist of precisely-defined biochemical steps which transform metabolites in an ordered fashion and with a predefined output (1). How these pathways were first assembled during evolution is not well understood (1), but they are known to have assembled at different evolutionary times (2–5), and subject to different selection pressures toward optimality (5–7) and robustness (8–10). Since metabolic innovation presented by a pathway is dependent on all enzymes functioning in a concerted manner, enzymes within a pathway are under selective pressure to become co-regulated (1,11–13). We therefore hypothesize that central metabolism has evolved with flux control mechanisms that are organized by pathway, to regulate their metabolic output.

Herein we test this hypothesis using the eukaryal model organism *Saccharomyces cerevisiae,* through a combination of chemostat cultivation, metabolic modeling, and mass spectrometry (MS)-based proteomics and phosphoproteomics. By correlating enzyme and phosphoenzyme levels with metabolic fluxes (14,15) across a wide range of conditions designed to perturb metabolism at the global scale (16), we identified reactions that are regulated by three mechanisms: (i) enzyme abundance, (ii) activating enzyme phosphorylation, and (iii) inactivating enzyme phosphorylation. We then tested whether these reactions are enriched in distinct pathways. We found that pathways in amino acid (AA) metabolism are predominantly regulated by enzyme abundance stemming from transcriptional control, whereas pathways in central carbon metabolism (CCM) and glycolipid/glycophospholipid (GL/GPL) metabolism are largely regulated by enzyme phosphorylation. In CCM, our dataset allowed us to map the function of 9 phosphoenzymes (12 phosphosites) which have not been mapped before; further, we observed a distinction in the function of enzyme phosphorylation (i.e. whether it activates or inactivates the enzyme) which coincides with the division between upper and lower glycolysis. In the GL/GPL pathway, we mapped the function of 14 phosphoenzymes (25 phosphosites) for the first time, and showed that the function of enzyme phosphorylation is coupled to the localization of enzymes in different cellular compartments. Finally, from an evolutionary perspective, we found that phosphoenzymes in central pathways arose in three distinct trajectories, largely coinciding with the division between CCM, GL/GPL metabolism, and AA metabolism. Taken together, our work reveals a non-random organization of flux control mechanisms related to enzyme abundance and phosphorylation in yeast central metabolic pathways, and provides an evolutionary history for the contribution of this organization to metabolic regulation.

## Results

### Data description

We previously described (16) a series of steady-state chemostat cultures of *S. cerevisiae,* designed to perturb cell metabolism at a global scale by modulating the cell growth rate and the central metabolism of carbon and nitrogen (Fig 1A). In this study, we performed Tandem Mass Tag (TMT)-based MS (17) to determine the protein and phosphoprotein abundances for 12 chemostat conditions in biological triplicates. A pooled sample was used as the internal reference in all TMT runs, which effectively means that the protein (P) and phosphoprotein (p) abundances were determined as the fold change relative to the mean of all samples (i.e. 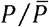 and 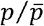, respectively). The phosphoprotein abundances were then normalized to the respective protein abundances, giving the relative fraction of each protein that has been phosphorylated 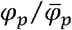; Table S1). The absolute abundance of proteins in these samples, in fmol per mg of cell dry weight (fmol/mgDW), was previously determined (16) by intensity-based absolute quantification (iBAQ), and can be found in Table S2.

**Figure 1.**
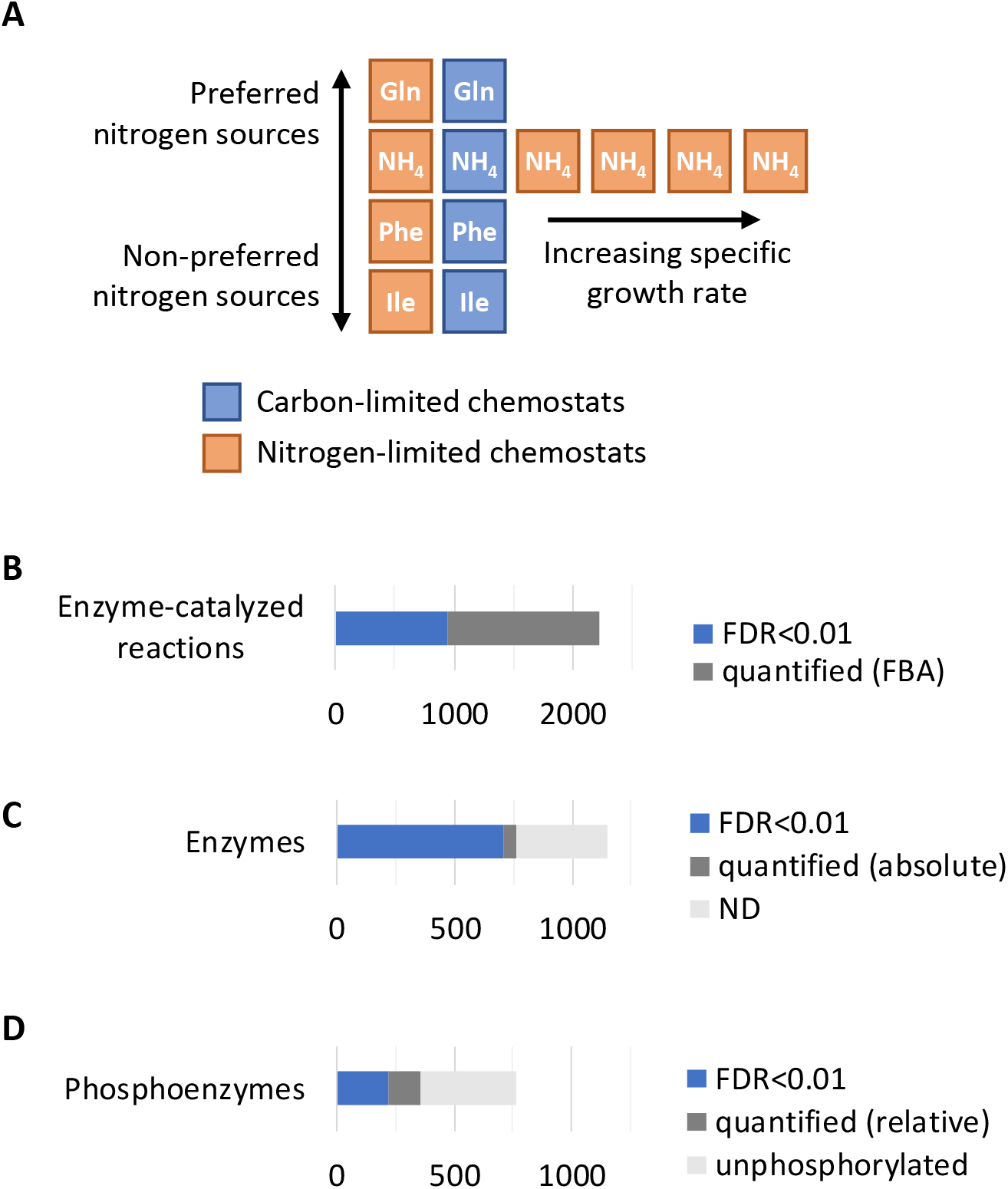
Regulation of metabolic flux by enzyme abundance and phosphorylation. (A) Experimental design to perturb cell metabolism at a global scale. Cells were grown in chemostats with different media compositions and at different growth rates. The nitrogen source of each chemostat condition is indicated. Colors represent either carbon-limited or nitrogenlimited conditions Full details can be found in supplementary file 1 of Yu *et al* (2021). (B) Number of enzyme-catalyzed reactions that carry non-0 flux in at least one condition, and those which carry significantly different flux (FDR<0.01 by 1-way ANOVA), as calculated by flux balance analysis (FBA) with random sampling. (C) Of all enzymes in yeast, the number of enzymes that are quantified (absolute quantification) and differentially expressed (FDR<0.01 by 1-way ANOVA) are shown. (D) Of all quantified enzymes, the number of phosphoenzymes that are quantified (relative quantification) and differentially expressed (FDR<0.01 by 1-way ANOVA) are shown.

To estimate metabolic flux, we performed flux balance analysis (FBA) (18) using the consensus yeast genome-scale metabolic model (GEM), Yeast8.4.0 (19), constraining the model using measured metabolite exchange fluxes and dilution rates (16). The feasible region of each chemostat condition was randomly sampled 1,000 times, allowing statistical significance to be calculated when fluxes were compared across conditions (20,21). The average flux for each reaction, calculated from the 1,000 random samplings, is in Table S3. This analysis showed that, out of 2,220 enzyme-catalyzed reactions that carry non-0 flux in at least one condition, 944 reactions (43%) carried significantly different flux in at least one condition (Fig 1B; FDR<0.01,1-way ANOVA), confirming that the different chemostat conditions have induced global changes in cell metabolism.

Out of a total of 1,148 enzymes in Yeast8.4.0, we detected 762 enzymes (66%) in our proteomics dataset (Table S2), of which 707 enzymes (93%) were differentially expressed (Fig 1C). In our phosphoproteomics dataset (Table S1), we found that 356 enzymes (47%) were phosphorylated at least once, in line with previous estimates (14,22). We further found that the 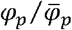 of 222 phosphoenzymes (62%) were significantly different in at least one condition (Fig 1D). Our dataset therefore represents the largest collection of steady-state flux-enzyme-phosphoenzyme relationships to date.

### Regulation of AA metabolism pathways by enzyme abundance

To identify reactions regulated by enzyme abundance, we calculated the correlation between metabolic flux and enzyme abundance for each reaction-enzyme pair wherein the reaction carries non-0 flux and the enzyme is detected in at least one condition, by Spearman correlation and maximal information coefficient (MIC) (23). The larger of the two calculated correlation scores was taken (Table S4), with a score of >0.5 as the cutoff. Of 2,123 reaction-enzyme pairs with calculated correlation scores, 371 (17%) were found to meet the cutoff. Enrichment of metabolic pathways was then performed using Generally Applicable Gene-set Enrichment (GAGE) (24,25). This revealed that reactions regulated by enzyme abundance are enriched primarily in pathways related to AA metabolism (Fig 2A). As an example, 8 out of 11 (73%) reaction-enzyme pairs in the lysine biosynthesis pathway are regulated by enzyme abundance, with the rest also clearly showing positive correlations between metabolic flux and enzyme abundance, although they did not meet the cutoff (Supplemental Fig 1 and Table S4; two examples are also shown in Fig 2B-C). Moreover, previous work has shown that for genes in AA metabolism, protein abundance is largely regulated by transcript abundance (16). Thus, flux through AA pathways is regulated by enzyme abundance (Fig 2A), which is, in turn, regulated transcriptionally, consistent with known transcriptional regulation programs in yeast nitrogen and amino acid metabolism (12,26).

**Figure 2.**
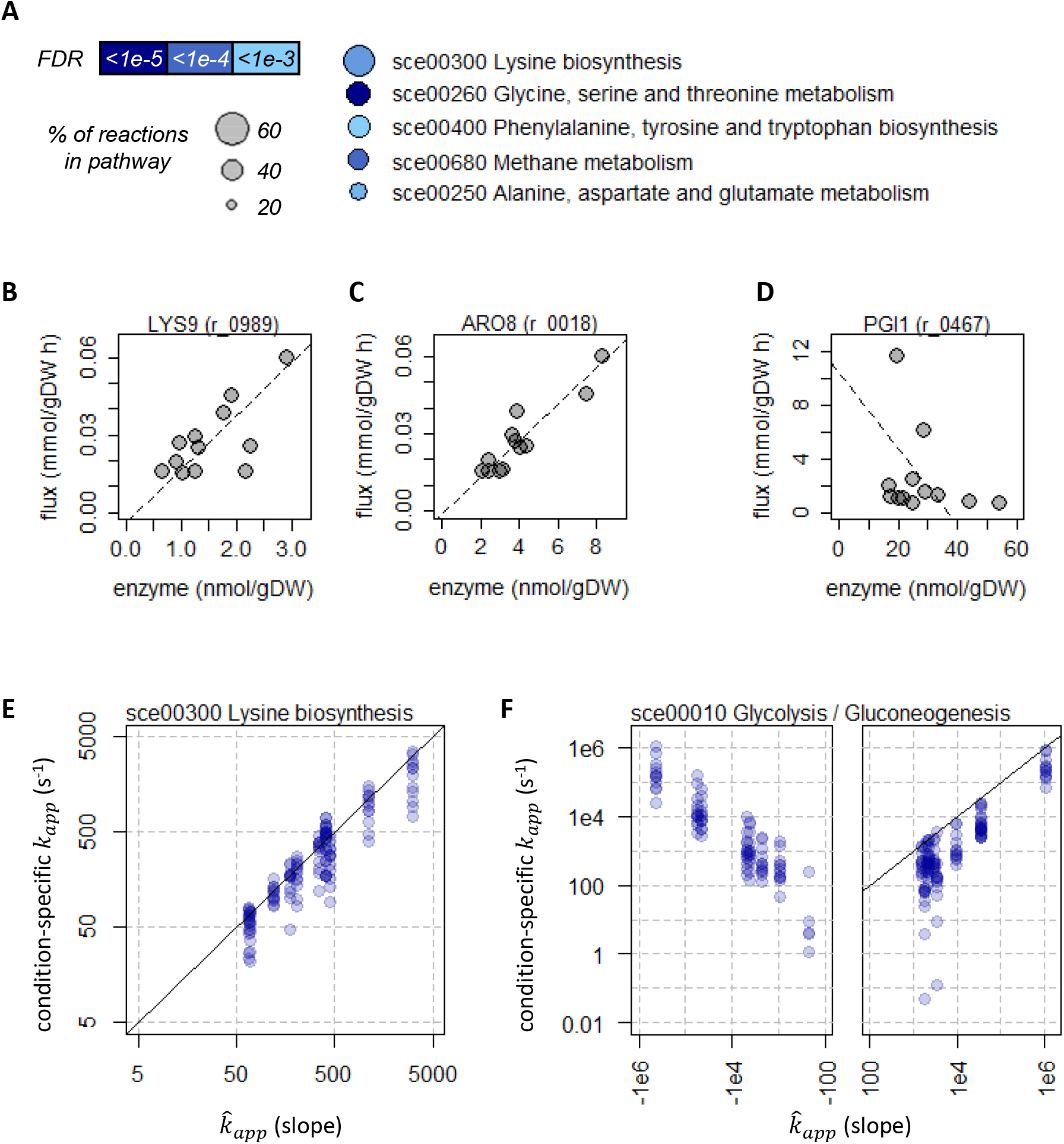
Amino acid metabolism pathways are regulated by enzyme abundance. (A) Reactions with high correlation between metabolic flux and enzyme abundance are enriched in metabolic pathways related to AA metabolism. The top 5 pathways with highest % of reactions in the pathway are shown. (B-C) Two example enzymes (reaction ID in brackets) in the lysine biosynthesis pathway showing high correlation between metabolic flux and enzyme abundance. The linear regression (dashed lines) are shown. (D) An example enzyme (reaction ID in brackets) in the glycolysis/gluconeogenesis pathway showing a paradoxical negative correlation between metabolic flux and enzyme abundance. The linear regression (dashed line) is shown. (E) Comparison between condition-specific *k_app_* (apparent catalytic rate) and 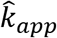 (slope of the linear regression between *v* and *E* of all conditions) in the lysine biosynthesis pathway. Solid black line is y=x. (F) Comparison between condition-specific *k_app_* values and 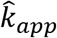 in glycolysis/gluconeogenesis. Solid black line is y=x.

Given the metabolic flux (*v*) and absolute-quantitative enzyme abundance (*E*), we calculated the apparent catalytic rate of the enzymes (*k_app_*) in each condition by

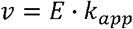

Additionally, we also calculated the slope of the linear regression between *v* and *E* of all conditions, designated 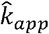. Note that although the value of 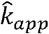 should be interpreted with caution, the comparison between 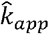 and condition-specific *k_app_* values is informative, particularly in cases where 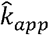 is negative, i.e. a lower enzyme abundance coincides paradoxically with a higher flux through the reaction. Such a relationship is prevalent in CCM (9,10), but cannot be captured in condition-specific *k_app_* values, since they are always positive; an example is given in Fig 2D. Here we found that, for pathways related to AA metabolism, 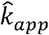 is nearly always positive, and agrees well with condition-specific *k_app_* values. Taking the lysine biosynthesis pathway as an example, we see that for each enzyme, 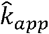 and condition-specific *k_app_* values generally agree to the order of magnitude (Fig 2E). We reasoned that these enzymes are likely operating near their turnover number (*k_cat_*) in all conditions tested, i.e. 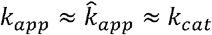, indicating that flux through these pathways are controlled predominantly by enzyme abundance, while other control mechanisms have little influence. Further, the 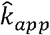 (and all *k_app_* values) for all 11 reaction-enzyme pairs in the pathway agree within two orders of magnitude (Fig 2E), consistent with the idea that the entire pathway is regulated by enzyme abundance to coordinate metabolic output.

We contrast these observations in AA metabolic pathways with a non-AA-pathway example, the glycolysis/gluconeogenesis pathway, shown in Fig 2F. In this pathway, a number of reaction-enzyme pairs have negative 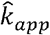 values (Fig 2F), consistent with previous reports (9,10) and directly demonstrating that enzyme abundance is not flux-controlling for these reactions. Moreover, the 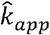 and condition-specific *k_app_* values differ by up to 5 orders of magnitude (Fig 2F), indicating that alternative mechanisms are in place to modulate flux depending on the growth condition. Taken together, these analyses show that pathways related to AA metabolism are coordinately regulated by enzyme abundance, while other pathways are regulated by other mechanisms.

### Regulation of CCM pathways by enzyme phosphorylation

We then performed phosphorylation regulation analysis (15), described by

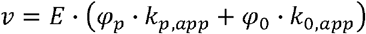

where *ϕ_p_* and *ϕ*_0_ are the fractions of phosphorylated and unphosphorylated enzymes, respectively; and *k_p,app_* and *k_0,app_* are the catalytic rate of phosphorylated and unphosphorylated enzymes, respectively. Of note, *k_p,app_* and *k_0,app_* are specific to each culture condition rather than being constant between conditions, similar to the way that *k_app_* of an enzyme is related to its *k_cat_* but is specific to each culture condition. By convention (15), we assume that if phosphorylation is functionally activating, then *k_p,app_ » k_O,app_*, which gives

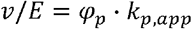

Since our phosphoproteomics data was measured as a relative fold change from the mean 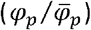, we transformed the enzyme-abundance-normalized flux (*v/E*) into a relative fold change from the mean as well (i.e. 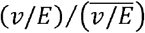; Table S5), which gives

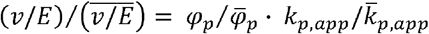

Here, since *ϕ_p_ + ϕ*_0_ = 1, a negative correlation between 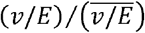 and 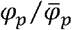 (relative flux decreases with increasing phosphoenzyme fraction) indicates that phosphorylation is functionally inactivating, i.e. *k_0,app_ » k_p,app_*. Wethen performed GAGE analysis (24,25), as before. This revealed that several pathways in CCM and GL/GPL metabolism contain a very high proportion of phosphoenzymes, which may be functionally activating, inactivating, or have no apparent effect in the conditions studied (Fig 3A).

**Figure 3.**
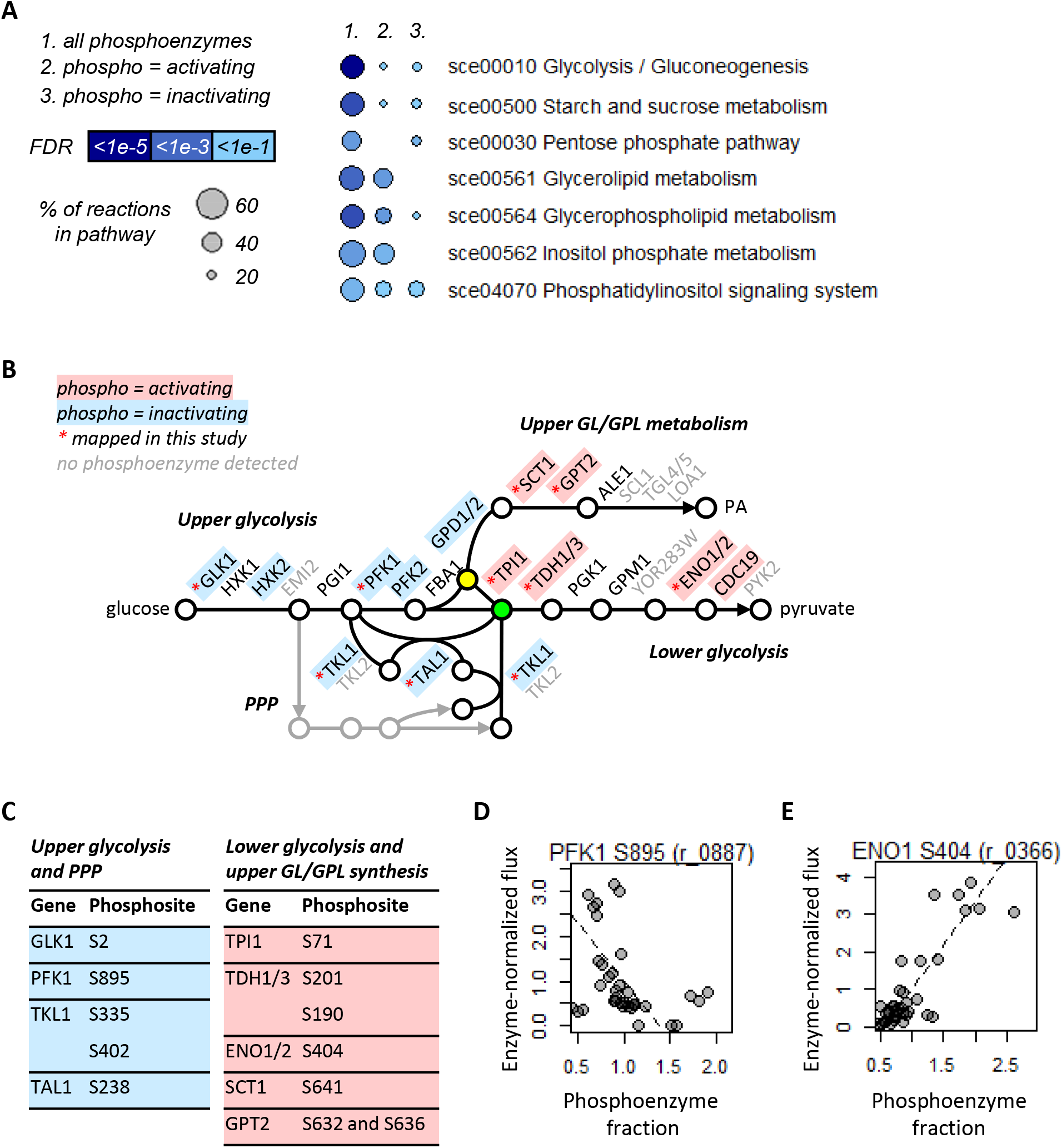
Central carbon metabolism pathways are regulated by enzyme phosphorylation. (A) Reactions for which enzymes are regulated by phosphorylation are enriched in metabolic pathways related to CCM and GL/GPL metabolism. Pathways which are enriched in group 1 and at least one of group 2 and 3 (phosphorylation activates or inactivates the enzymes in the pathways, respectively), are shown. (B) Schematics of glycolysis, pentose phosphate pathway (PPP), and upper glycerolipid/ glycerophospholipid (GL/GPL) metabolism showing phosphoenzymes that are activated (pink) or inactivated by phosphorylation (blue). Phosphoenzymes that are functionally mapped for the first time in this study are indicated by red asterisks. The metabolites dihydroxyacetone phosphate (DHAP) and glyceraldehyde 3-phosphate (G3P) are marked in yellow and green, respectively. (C) Phosphosites of phosphoenzymes that are functionally mapped in this study. Colors are as in (B). (D-E) Two example phosphoenzymes (reaction ID in brackets) showing a negative (D) and a positive (E) correlation between metabolic flux and phosphoenzyme fraction. The linear regression (dashed lines) are shown.

We first focused on the pathways of glycolysis/gluconeogenesis, pentose phosphate pathway (PPP), and GL/GPL metabolism, which branches off from glycolysis at dihydroxyacetone phosphate (DHAP; Fig 3B). Many enzymes in these pathways have previously been found to be phosphorylated in high-throughput phosphoproteomics studies, but whether these phosphorylation events carry function (i.e. associated with increased or decreased flux through the reaction) has only been mapped for a few enzymes (22,27). Moreover, for the few phosphoenzymes that has been functionally mapped, some enzymes are activated by phosphorylation (e.g. CDC19 at S22 (28)), while others are inactivated (e.g. HXK2 at S158 (29); PFK2 at S163 (14)). Thus it seemed that flux control by enzyme phosphorylation operated on an enzyme-specific basis, although data has been too sparce to make this claim conclusively. Here in our dataset, in addition to the previously identified functional phosphoenzymes/phosphosites, we further identified 4 phosphoenzymes (5 phosphosites) that are inactivated by phosphorylation, and 5 phosphoenzymes (7 phosphosites) that are activated in these pathways (Fig 3C; two examples are shown in Fig 3D-E). Importantly, we observed a pathway-level division between phosphoenzymes that are activated or inactivated by phosphorylation, which coincides with the division between upper and lower glycolysis, separating the metabolism of 6-carbon molecules and 3-carbon molecules at the step of glyceraldehyde 3-phosphate (G3P) and DHAP production (Fig 3B-C). Thus, flux control by enzyme phosphorylation is indeed coordinated at the pathway level in CCM, rather than operating on an enzyme-specific basis.

### Regulation of lower GL/GPL metabolism by enzyme phosphorylation

We next examined the regulation of lower GL/GPL metabolism, which extends from phosphatidic acid (PA) into several large branches to produce (and degrade) diacylglycerol (DAG) and triacylglycerol (TAG); phosphatidylserine (PS), phosphatidylethanolamine (PE), and phosphatidylcholine (PC); as well as phosphatidylinositol (PI) and PI derivatives (Fig 4A). Similar to enzymes in CCM, several enzymes in lower GL/GPL metabolism are known to be phosphorylated, but functional mapping has been restricted to a few cases (e.g. multiple sites on CDS1 (30), PAH1 (31), and CHO1 (30,32)), with a mix of activating and inactivating functions. Here, we report the functional mapping of 14 enzymes (25 phosphosites) in GL/GPL metabolism, showing for the first time that nearly every reaction in this pathway is regulated by enzyme phosphorylation (Fig 4B). In terms of function, however, we found a mix of activating and inactivating phosphorylation events on different enzymes, and in a few cases on different phosphosites of the same enzyme as well (ARE2, PAH1, PIK1, and FAB1; Fig 4A-B). This complexity suggests that additional factors may have co-evolved with enzyme phosphorylation to regulate flux through this pathway in a coordinated manner. Indeed, we observed that the function of enzyme phosphorylation is coupled to the localization of the enzymes, with enzymes localized in the endoplasmic reticulum (ER) being predominantly inactivated by phosphorylation, while others are predominantly activated (Fig 4B). Thus, the GL/GPL metabolic pathway is regulated by a combination of enzyme phosphorylation and metabolic compartmentalization.

**Figure 4.**
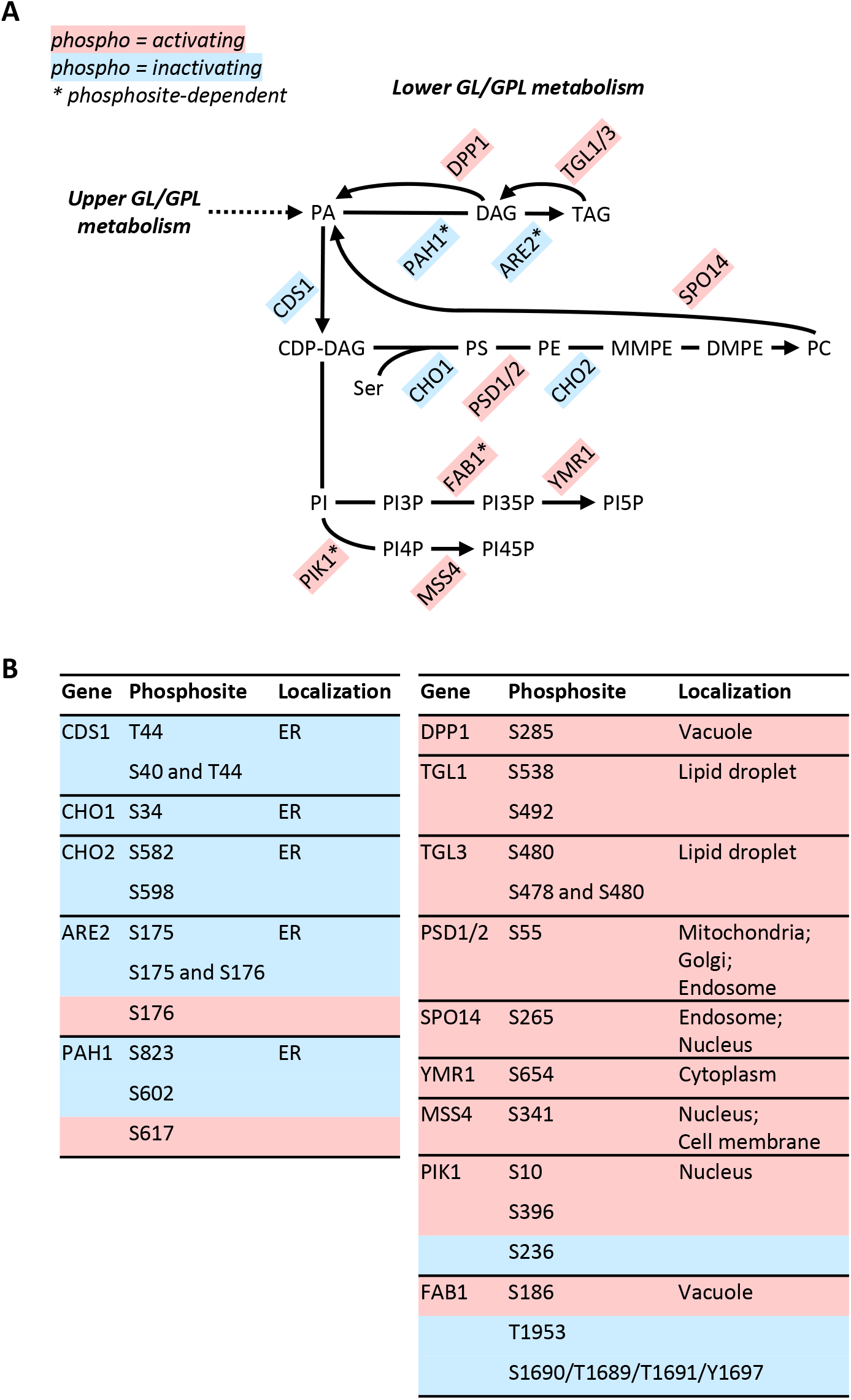
Lower glycerolipid/glycerophospholipid pathways are regulated by enzyme phosphorylation. (A) Schematics of lower glycerolipid/glycerophospholipid (GL/GPL) metabolism showing phosphoenzymes that are activated (pink) or inactivated by phosphorylation (blue). Phosphoenzymes with phosphosite-dependent activities are marked with asterisks. Enzymes with phosphosites that are functionally mapped for the first time in this study are shown. PA, phosphatidic acid; DAG, diacylglycerol; TAG, triacylglycerol; CDP-DAG, cytidine diphosphate-diacylglycerol; PS, phosphatidylserine; PE, phosphatidylethanolamine; MMPE, monomethyl-phosphatidylethanolamine; DMPE, dimethyl-phosphatidylethanolamine; PC, phosphatidylcholine; PI, phosphatidylinositol; PI(x)P, phosphatidylinositol (x)-phosphate. (B) Phosphosites of phosphoenzymes that are functionally mapped in this study. Colors are as in (A). Localization of the enzymes are indicated. ER, endoplasmic reticulum.

### Evolution of phosphorylation as a regulatory mechanism in central metabolism

To understand the evolutionary history of enzyme phosphorylation as a pathway-level flux control mechanism, we reconstructed the trajectory of phosphoenzyme evolution in central pathways based on the estimated age of fungal phosphosites on metabolic enzymes (33). We first focused on CCM (Fig 5A) and GL/GPL metabolism (Fig 5B), as we have shown coordinated flux regulation by enzyme phosphorylation in these pathways. In CCM pathways, we found that the proportion of enzymes with conserved phosphosites increased gradually from 0-20% at ~700 million years ago, to 50-60% in the present day (Fig 5A). As a comparison, CCM pathways are evolutionarily ancient: for example, lower glycolysis has been conserved since the last universal common ancestor (LUCA) ~3.5 billion years ago (1,4). Thus, the evolution of phosphorylation as a regulatory mechanism in CCM is a relatively late adaptation, occurring after the divergence of Eukarya from Archaea and the subsequent expansion of protein phosphorylation in Eukarya (34,35). GL/GPL metabolism, on the other hand, is a much younger pathway in Eukarya, with its basic biochemical structures being formed after the emergence of the last eukaryal common ancestor (LECA) ~2.1 billion years ago (3,36). Genes in this pathway, however, continued to diverge, leading to the evolution of young (between 900 and 400 million years old) and highly lineage-specific enzymes (3). Consistently, we found that GL/GPL pathway enzymes with conserved phosphosites first emerged between 400 and 200 million years ago, which rapidly increased to encompass 70-100% of enzymes in this pathway in the present day (Fig 5B). This coincides with the emergence of the budding yeast common ancestor (BYCA) ~400 million years ago (37), suggesting that an early selection pressure in the budding yeasts had favoured the co-evolution of enzymes in the GL/GPL pathway and their coordinated regulation by phosphorylation.

**Figure 5.**
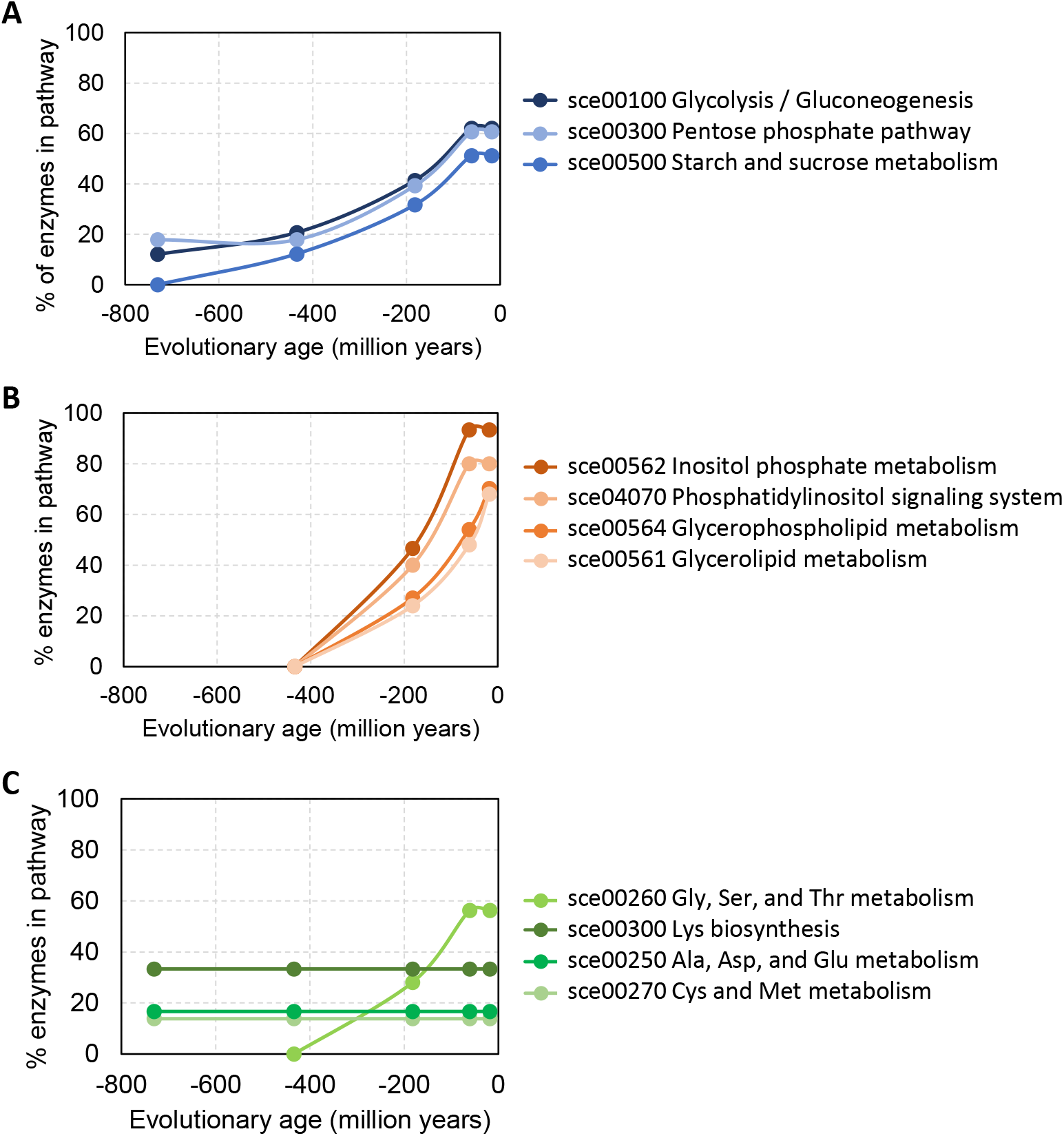
Evolutionary history of enzyme phosphorylation in central pathways, based on the estimated age of fungal phosphosites on metabolic enzymes in Studer *et al,* 2016 (reference 33). (A) The % of enzymes with conserved phosphosites in CCM pathways as a function of evolutionary time. (B) The % of enzymes with conserved phosphosites in GL/GPL metabolism pathways as a function of evolutionary time. (C) The % of enzymes with conserved phosphosites in AA metabolism pathways as a function of evolutionary time.

Interestingly, we also observed a third distinct trajectory of phosphoenzyme evolution in several AA metabolism pathways (Lys biosynthesis; Ala/Asp/Glu metabolism; and Cys/Met metabolism), though not all (Gly/Ser/Thr metabolism; Fig 5C). In this trajectory, a constant fraction (20-40%) of enzymes contain phosphosites that have been conserved since >700 million years ago (Fig 5C). Typically, such a strong conservation is indicative of functional importance (38,39); yet in our dataset, these phosphorylation events are generally not correlated with flux (Fig 3A, Table S6). On the other hand, previous work has shown that more conserved phosphosites are more likely to contribute to the stability of protein-protein interactions (33), pointing to the possibility that these phosphosites facilitate the interaction between enzymes to form enzyme complexes, which are known to be influenced by enzyme phosphorylation (40–42). Finally, we note that Gly/Ser/Thr metabolism exhibits a phosphoenzyme evolution trajectory (Fig 5C) similar to that of GL/GPL metabolism (Fig 5B), suggesting that Gly/Ser/Thr metabolism were under selection pressure to become co-regulated with GL/GPL pathways in the evolution of budding yeasts, in order to supply Ser for PS synthesis (Fig 4A). Our analyses therefore demonstrate that distinct trajectories of evolutionary history gave rise to the pathway-specific flux control mechanisms in CCM, GL/GPL metabolism, and AA metabolism.

## Discussion

Understanding the regulation of metabolism is a key goal in biology. Here we reveal that flux control mechanisms in yeast central metabolism are organized by pathway, wherein (i) AA metabolism are predominantly regulated by enzyme abundance stemming from transcriptional regulation; (ii) upper glycolysis and PPP, by inactivating enzyme phosphorylation; (iii) lower glycolysis with the branching off upper GL/GPL metabolism, by activating enzyme phosphorylation; and (iv) lower GL/GPL metabolism, by a combination of enzyme phosphorylation and metabolic compartmentalization. We further delineate the evolutionary history of the organization of flux control mechanisms in yeast central metabolism, showing distinct evolutionary trajectories of phosphoenzymes within CCM, GL/GPL metabolism, and AA metabolism. Our work highlights that the complex regulation of metabolism contains organizational structures that have been shaped by evolutionary selection, and warrants the examination of additional flux control mechanisms to further our understanding of both metabolic regulation and its evolution.

This study reports the largest collection of flux-enzyme-phosphoenzyme relationships to date, in the eukaryal model organism *Saccharomyces cerevisiae.* Importantly, as the experimental conditions were designed to perturb cell metabolism at a global scale, 43% of metabolic reactions in this dataset carry significantly different flux; 93% of enzymes are differentially expressed; and 62% of phosphoenzymes are differentially phosphorylated (Fig 1C-D). Our dataset is therefore particularly suitable to investigate metabolic regulation by enzyme abundance and phosphorylation. This allowed us to map the function of 9 phosphoenzymes (12 phosphosites) in CCM (Fig 3C), and 14 phosphoenzymes (25 phosphosites) in GL/GPL metabolism (Fig 4B), which have not been reported before. Moreover, the use of steady-state chemostat conditions in this dataset removes the confounding factors related to time (e.g. propagation of regulatory signals), which simplifies parameter identification and facilitates the discovery of regulatory interactions. The caveat with this is that regulatory interactions that are transient in nature are not captured. This is unlikely to influence the effects related to enzyme abundance, but metabolic regulation by enzyme phosphorylation could be more prevalent than reported here (12% of reaction-phosphoenzyme pairs; Table S6), which remains to be discovered.

Co-regulation of enzymes in the same pathway is an intuitive way to regulate metabolism, and indeed it is well-established in transcriptional regulation circuitries such as bacterial operons (43) and eukaryal regulons (44,45). In yeast, genes related to nitrogen and amino acid metabolism are among the best known to be transcriptionally co-regulated, in the nitrogen catabolite repression (NCR) and general amino acid control (GAAC) regulons (12,26). Consistently, our results show that pathways in AA metabolism are predominantly regulated by enzyme abundance (Fig 2A) stemming from transcriptional regulation (16). A rational extension of these observations is the hypothesis that post-transcriptional control mechanisms may be similarly coordinated by pathway, which is supported by previous reports of coordinated protein translation (9) as well as transcript and protein turnover rates (46) in central metabolic pathways. Our results further support this hypothesis, demonstrating that flux control by enzyme phosphorylation in CCM and GL/GPL metabolism are also organized at the pathway level (Fig 3B and Fig 4A). In contrast, substrate-level or allosteric enzyme inhibition do not appear to be enriched in particular pathways (47), highlighting that multiple organizational structures can be present in metabolic regulation.

In CCM, we found a divide between the function of enzyme phosphorylation coinciding with the division between upper and lower glycolysis, i.e. between the metabolism of 6-carbon molecules and 3-carbon molecules (Fig 3B). As glycolysis is among the most ancient and conserved pathways in life (1,2), this appears to be contradictory to the idea of co-evolution between a pathway and its regulatory mechanism(s). However, whereas lower glycolysis is near-universal in life (4,48), upper glycolysis exists in several distinct variants, most notably the Embden-Meyerhof-Parnas (EMP) pathway present in most organisms, and the Entner-Doudoroff (ED) pathway in certain bacteria and archaea (4,7,49). Moreover, within the EMP pathway, nearly all enzymes are derived from multiple gene sequence families (4), suggesting that they have independently evolved several times. Thus, while the biochemistry of glycolysis is ancient and conserved, the selection pressures on glycolytic enzymes were clearly distinct between upper and lower glycolysis: enzymes in lower glycolysis were strongly conserved against replacement and mutations (50), while enzymes in upper glycolysis were lost multiple times and re-gained through recruitment or lateral gene transfer (4). As such, it is probable that different regulatory signals have co-evolved with enzymes in upper and lower glycolysis, as we have shown here.

In GL/GPL metabolism, we found that flux through this pathway is regulated by a combination of enzyme phosphorylation and metabolic compartmentalization (Fig 4B). Independently, both mechanisms are known to play key roles in metabolic regulation (13,22); in particular, the spatial organization of metabolic reactions in different subcellular compartments can influence both enzyme efficiency (e.g. by providing different microenvironments) and pathway efficiency (e.g. by channelling metabolites), and is a hallmark of eukaryal cells. Here we showed that, when combined, these two regulatory mechanisms can give rise to an unexpected complexity in the regulation of GL/GPL metabolism, wherein the function of enzyme phosphorylation is coupled to the subcellular compartment. These results illustrate that complex interactions can arise when multiple regulatory mechanisms co-evolve with a metabolic pathway, particularly for relatively young pathways evolving in a relatively complex cell system such as eukaryal cells.

Lastly, we note that while a number of enzymes in AA metabolism are phosphorylated, their function could not be mapped as the phosphoenzyme abundances were not correlated with reaction fluxes (Fig 5C and Table S6). This is consistent with a previous study showing that phosphorylation of 2 enzymes (5 phosphosites) in the lysine biosynthesis pathway, either singly or in combination, have little biological consequence (27). On the other hand, the relatively old evolutionary age and high degree of conservation of phosphosites in AA metabolism enzymes (Fig 5C) suggest that they carry some function. Since more conserved phosphosites are more likely to contribute to the stability of protein-protein interactions (33), this points to the possibility that phosphorylation of these enzymes functions to facilitate protein-protein interactions, rather than directly influencing enzyme activity. Close interaction of enzymes mediated by phosphorylation has been previously described for several pathways (40–42), with the most notable example being enzymes in *de novo* purine biosynthesis, which assemble into macromolecular complexes called purinosomes and is regulated by phosphorylation of key enzymes (40). It will be interesting to test whether the highly-conserved phosphosites in AA metabolism enzymes function in a similar capacity.

In conclusion, we demonstrated that yeast central metabolism exhibits a non-random, pathway-level organization of flux control mechanisms related to enzyme abundance and phosphorylation. Delineating the organizational structures of additional flux control mechanisms, as well as their evolutionary histories, will have important implications for bioengineering and medicine.

## Materials and methods

### Culture conditions

Culture conditions are as previously described (16). The yeast *Saccharomyces cerevisiae* CEN.PK113-7D (MATa, MAL2-8c, *SUC2)* was used for all experiments. Cells were stored in aliquoted glycerol stocks at −80 °C. Chemostat experiments were carried out in DASGIP 1L bioreactors (Jülich, Germany) equipped with off-gas analysis, pH, temperature and dissolved oxygen sensors. Chemostat experiments were carried out at 30 °C, pH 5, working volume 0.5 L, aeration 1 vvm, pO_2_ >30%, agitation speed 800 rpm. Additional media components are: KH_2_PO_4_, 3 g L^-1^; MgSO_4_·7H_2_O, 0.5 g L^-1^; trace metals solution, 1 ml L^-1^; vitamin solution, 1 ml L^-1^; antifoam, 0.1 ml L^-1^. The trace metals solution contained: EDTA (sodium salt), 15.0 g L^-1^; ZnSO_4_·7H_2_O, 4.5 g L^-1^; MnCI_2_·2H_2_O, 0.84 g L^-1^; COCI_2_·6H_2_O, 0.3 g L^-1^; CuSO_4_·5H_2_O, 0.3 g L^-1^; Na_2_M·O_4_·2H_2_O, 0.4 g L^-1^; CaCI_2_·2H_2_O, 4.5 g L^-1^; FeSO_4_·7H_2_O, 3.0 g L^-1^; H_3_BO_3_, 1.0 g L^-1^; and KI, 0.10 g L^-1^. The vitamin solution contained: biotin, 0.05 g L^-1^; p-amino benzoic acid, 0.2 g L^-1^; nicotinic acid, 1 g L^-1^; Ca-pantothenate, 1 g L^-1^; pyridoxine-HCl, 1 g L^-1^; thiamine-HCl, 1 g L^-1^ and myo-inositol, 25 g L^1^.

### Sampling from bioreactor

The sampling procedure for proteome and phosphoproteome measurements was as previously described (16). Briefly, the reactor dead volume was collected with a syringe and discarded. Biomass was collected from the reactor with a syringe and injected into 50 ml Falcon tubes chilled on ice. Samples were centrifuged for 4 min at 3,000 x g at 4 °C; cell pellets were washed once with 20 ml of chilled dH_2_O, washed again with 1 ml of chilled water, transferred into Eppendorf tubes, flash frozen in liquid nitrogen, and stored at −80 °C until analysis.

### Quantitative proteome and phosphoproteome measurements

Sample preparation for proteomic analysis was performed as reported in our previous publication (16) with a few substantial differences as outlined below.

Pierce Phosphatase Inhibitor tablet (A32957; Thermo Fischer Scientific, Waltham, MA) was added to the lysis buffer, 800 μg of total protein from each lysate was digested and labeled with TMT 10plex isobaric reagents, ca. 10% of the amount from each combined TMT-labeled set was reserved for the total protein analysis, while ca. 90% of the amount was reserved for phosphopeptide enrichment. Each combined TMT-set was separated into 20 primary fractions using basic-pH reversed-phase (bRP)-HPLC, which were pooled into 10 final fractions for analysis. All fractions for the non-enriched and phosphopeptide TMT-sets were analyzed using an MS2-TMT method (HCD energy 38%, MS2 resolution 50,000) on an Easy-nLC coupled to an Orbitrap Fusion Lumos mass spectrometer (both Thermo Fischer Scientific); each non-enriched fraction was analyzed over a 90 min LC gradient.

The remaining amount of ca. 90% for each combined TMT-set was purified on Pierce peptide desalting spin columns (Thermo Fisher Scientific) according to the manufacturer’s instructions and separated into 2 equal aliquots. One aliquot per TMT set was used for the phosphopeptide enrichment on High-Select™ Fe-NTA phosphopeptide enrichment kit, another aliquot for each TMT set was processed using the High-Select™ TiO_2_ phosphopeptide enrichment kit (both Thermo Fisher Scientific), according to the manufacturer’s protocols. Eluates were immediately put into Speedvac and evaporated at room temperature until the volumes have decreased ca. 2-3 times. Enriched phosphopeptides from TiO_2_ and FeNTA were combined for each TMT set and separated into 20 primary/10 final fractions using bRP-HPLC. Each final phopshopeptide fraction was analyzed using an MS2-TMT method over a 120 min LC gradient.

LC-MS data was processed via Proteome Discoverer version 2.4 (Thermo Fisher Scientific). The files were matched by Mascot 2.5.1 (Matrix Science, London, UK) against the reference *Saccharomyces cerevisiae* ATCC 204508 / S288c database from Uniprot (November 2019), supplemented with common proteomic contaminants, with trypsin (max. 1 missed cleavage) as an enzyme rule, precursor ion tolerance of 10 ppm and fragment ion tolerance of 0.03 Da; TMT 6-plex on lysine and peptide N-termini, methylthiolation on cysteine were set as fixed modifications; oxidation on methionine was set as a variable modification. Additionally, phosphorylation on S, T and Y was set as a variable modification for the enriched samples. Percolator node was used for PSM validation with the strict FDR threshold of 1%. IMP-ptmRS node was used to assess the site localization probabilities.

Raw LC-MS data, result files and the comprehensive description of sample preparation and data processing have been uploaded to the PRIDE (51) repository with the dataset identifier PXD029709.

### Data processing and analysis

For flux balance analysis (FBA), MATLAB R2019b (MathWorks, Inc., Natick, MA) with Gurobi solver (Gurobi Optimizer, Beaverton, OR) in the COBRA toolbox (52) was used for flux balance analysis. The consensus yeast metabolic model Yeast8.4.0 (https://github.com/SysBioChalmers/yeast-GEM) (19) was constrained by the nutrient and waste product exchange fluxes, and the specific growth rate, of each condition (16). Condition-dependent biomass composition was introduced by scaling the coefficients of the protein and RNA pseudo-reactions to equal to measured protein and RNA abundance, as previously performed (9). The constrained model for each condition was then randomly sampled (20) 1,000 times wherein random pairs of reactions are selected as objective functions and assigned at random weights (20), as implemented in the RAVEN toolbox (21). The average flux for each reaction is calculated and can be found in Table S3. Reactions that carry non-0 flux in all random samplings is then compared by 1-way ANOVA with a FDR<0.01 cutoff.

For phosphoproteomics data, phosphopeptides that could be mapped to multiple proteins were reported as a phosphopeptide of each protein, with indication that it is a multi-mapped peptide (value of 1 in the “multi_mapped” column of Table S1). Phosphopeptides that were detected in at least 50% of all samples, and that were detected in at least one biological replicate in each experimental condition, were included for further analysis. For normalization of phosphopeptide abundance against total protein abundance, first the mean of protein abundance for the three biological replicates of each condition was taken. Proteins that were detected in at least one replicate in all conditions were included for further analysis. Phosphopeptide abundance of each sample was then normalized against the mean total protein abundance of the three biological replicates of each condition. Normalized phosphopeptide abundance is given in Table S1.

Correlation between metabolic flux and enzyme abundance for each reaction-enzyme pair, and between enzyme-abundance-normalized metabolic flux and phosphoenzyme fraction, was calculated by Spearman correlation and maximal information coefficient (MIC) (23). The larger of the two calculated correlation scores was taken (Table S4), with a score of >0.5 as the cutoff. Enzyme regulation by phosphorylation was analyzed as previously described (15). For metabolic pathway enrichment of abundance-regulated and/or phosphorylation-regulated enzymes, Generally Applicable Gene-set Enrichment (GAGE) was performed (24,25) with enzyme names as the input. Localization of enzymes in the GL/GPL metabolic pathway were collected from literature (53,54).

Reconstruction of the evolutionary history of enzyme phosphorylation at the pathway level is performed using the estimated age of fungal phosphosites (33) of on enzymes in *S. cerevisiae,* found in YeastGEM8.4.0 (19). Enzymes with phosphosites estimated to have evolved at Y0 (731 Mya), Y1 (434 Mya), Y2-Y6 (182-61 Mya), and S1-Sc (18 Mya and younger) (33) were subject to GAGE analysis for metabolic pathway enrichment (24,25). For simplicity, enzymes with multiple phosphosites which have different estimated evolution times were excluded. The % of enzymes in the pathway which have evolved phosphosites at each time point is calculated as the cumulative sum until that time point.

### Data and code availability

The mass spectrometry proteomics data have been deposited to the ProteomeXchange Consortium via the PRIDE (51) partner repository with the dataset identifier PXD029709. Computer scripts (Matlab and R) used for this study are submitted with the present work as computer_code_20210721.zip. Processed proteomics data are in Table S1-2. FBA with random sampling results are in Table S3. Table S4-6 contain all numerical data used to produce figures.

## Acknowledgements

We thank members of the Nielsen lab for discussions.. This research was supported by funding from the Novo Nordisk Foundation (grant number NNF10CC1016517) and the Knut and Alice Wallenberg Foundation.

---

**Supplemental Figure 1, related to Figure 2.**
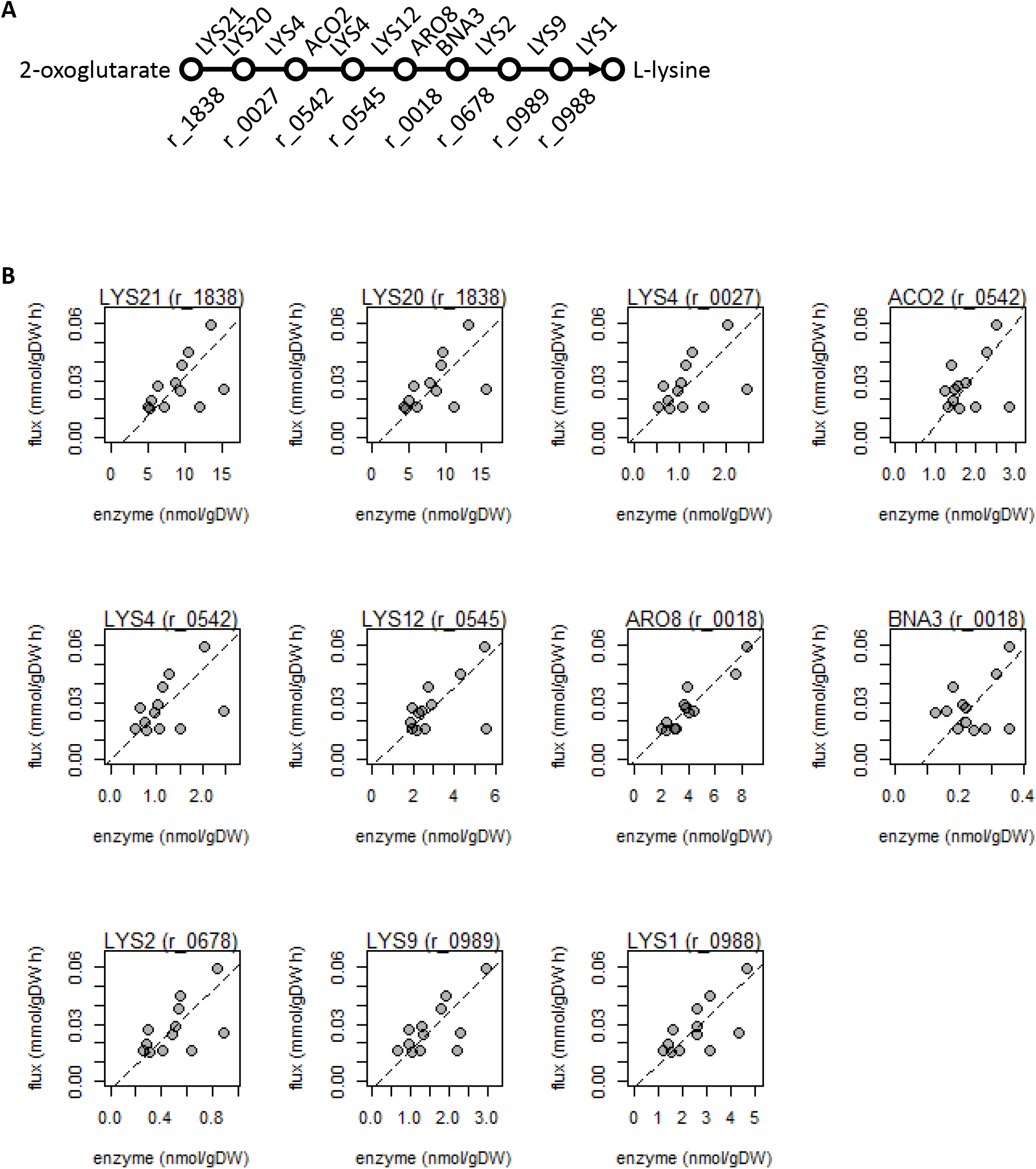
Lysine biosynthesis is regulated by enzyme abundance. (A) Schematics of the lysine biosynthesis pathway showing enzymes and reaction IDs of each reaction. (B) All enzymes (reaction ID in brackets) in the lysine biosynthesis pathway showing high correlation between metabolic flux and enzyme abundance. The linear regression (dashed lines) are shown.

